# Collagen targeting IL-12 combined with Doxorubicin enhances the anti-tumor effect against osteosarcoma

**DOI:** 10.64898/2026.05.07.723520

**Authors:** Takumi Matsuo, Laurine Noblecourt, Pooja Kaur, Chuyi Wang, Po-Chuan Chiu, Koichi Sasaki, Charanjit Singh, Adrian Larkeryd, Anguraj Sadanandam, Paul H. Huang, Jun Ishihara

## Abstract

Osteosarcoma (OS) is the most prevalent primary bone malignancy in children and adolescents; however, therapeutic outcomes remain suboptimal due to tumor heterogeneity, chemoresistance, and inadequate immune activation. Doxorubicin (Dox), the standard therapy that induces immunogenic cell death, has its efficacy compromised by the immunosuppressive tumor microenvironment (TME). While interleukin-12 (IL-12) can activate and recruit various immune cells, making it an attractive combination partner, its systemic delivery is severely limited by dose-limiting toxicity. We have previously reported that intravenous injection of A3 collagen binding domain (CBD) of von Willebrand Factor preferentially accumulates into the TME of various tumor models enriched in collagen I and III. Furthermore, CBD-fused IL-12 (CBD-IL-12) demonstrated superior therapeutic effects against various cancer models compared to unmodified IL-12 due to its collagen-targeted delivery and the resulting tumor-localized inflammation. Given that the OS TME also exhibits higher collagen I and III expression compared to normal bone, we hypothesized that a CBD-IL-12 fusion protein could showcase potent anti-tumor efficacy in OS via tumor-specific accumulation. Here, we demonstrated that CBD-IL-12 exhibited 4-fold enhanced tumor accumulation compared to unmodified IL-12 and increased cytotoxic T cell infiltration by 2.2-fold within the immune-cold microenvironment in a mouse model of OS. The combination of CBD-IL-12 with Dox significantly prolonged median survival in two independent murine OS models. This coordinated approach utilizing Dox coupled with precision-targeted IL-12 immunotherapy represents a clinically translatable strategy that overcomes the inherent limitations of single-agent treatments for OS.

**Highlight:**

- Collagen-targeted IL-12 increases tumor accumulation in osteosarcoma.
- The collagen-targeted IL-12 synergizes with doxorubicin in osteosarcoma models.
- Combination therapy enhances T cell differentiation and activates innate immunity.

## Introduction

Osteosarcoma (OS) represents the most common primary bone malignancy in children and adolescents, accounting for approximately 60% of all bone sarcomas in patients under 25 years of age[1–3]. Despite significant advances in multimodal therapy, the overall survival rate for patients with localized disease is 70-75 %, while that for patients with metastatic OS is 20-30 %[4]. The standard of care relies on surgery with neoadjuvant or adjuvant chemotherapeutics, including doxorubicin (Dox), cisplatin, and high-dose methotrexate, yet treatment resistance and severe systemic toxicity limit therapeutic efficacy and patient quality of life[5, 6]. Dox, a cornerstone of OS chemotherapy, exerts its cytotoxic effects through DNA intercalation, topoisomerase II inhibition, and generation of reactive oxygen species generation[7, 8]. Critically, Dox induces immunogenic cell death (ICD), releasing damage-associated molecular patterns (DAMPs) that recruit dendritic cells (DCs) and enhance its maturation[9, 10]. This could subsequently trigger T cell priming, creating temporal windows for synergistic immunotherapy combinations, particularly with cytokines that amplify antigen-specific T cell responses[11, 12].

Cancer immunotherapy has emerged as a promising approach for the patients with various types of cancer. Although checkpoint-inhibitor (CPI) therapy achieved clinical success[13], most patients with established tumors demonstrate limited responses due to the lack of immune infiltration in the tumor microenvironments (TME)[14]. The TME in OS typically exhibits lower immune cell infiltration patterns, characterized by sparse T cell infiltration and predominant immunosuppressive myeloid populations, rendering them unresponsive to CPI therapies[15, 16]. This immune evasion is further compounded by the mineralized extracellular matrix (ECM) environment, which creates physical barriers to immune cell infiltration and therapeutic drug delivery[17, 18]. Consequently, there exists an urgent clinical need for combination strategies that simultaneously overcome physical delivery barriers while enhancing immune activation within the TME[19].

Interleukin-12 (IL-12), which is a cytokine composed of p40 and p35 subunits, reshape immunosuppressive TME through Interferon-γ (IFN-γ)-mediated immune activation[20, 21]. IL-12 stimulates cytotoxic T cells and promotes Th1 polarization, directly counteracting immunosuppressive mechanisms[22]. Multiple preclinical studies have demonstrated striking therapeutic effects against various tumor models[23, 24]. The therapeutic potential of IL-12 has been also validated in preclinical OS models. Local IL-12 delivery via viral vectors reduced tumor burden and pulmonary metastasis[25]. Engineered T cells expressing IL-12 demonstrated substantial therapeutic effect against different patient-derived xenograft models of OS[26]. Furthermore, preclinical studies demonstrated that IL-12 therapy synergizes with chemotherapy through complementary mechanisms: chemotherapy can induce ICD, releasing tumor antigens and danger signals, while IL-12 enhances DC activation and antigen cross-presentation, thereby amplifying tumor-specific T cell priming and cytotoxic responses[27]. However, systemic IL-12 administration produces dose-limiting immune-related adverse events (irAEs), severely restricting its therapeutic window and clinical utility[28–30].

Tumor-targeted delivery systems have emerged to harness IL-12’s therapeutic potential while minimizing systemic toxicities, either through intratumoral delivery or engineering IL-12 fusion proteins to exploit unique features of TME[31, 32]. We previously demonstrated that the A3 collagen-binding domain (CBD) of von Willebrand factor (vWF) effectively accumulates within various types of tumors by targeting exposed collagen I and III, due to the more permissive tumor vasculature[33]. IL-12 fused with the CBD (CBD-IL-12) exhibited enhanced tumor accumulation following systemic administration without unfavorable localization to healthy tissue in comparison to unmodified IL-12. This resulted in the sustained intratumoral levels of IFN-γ, while concurrently leading to a substantial reduction in systemic levels of IFN-γ and organ damage, in contrast to unmodified IL-12. Consequently, CBD fusion to IL-12 enhanced both safety and therapeutic efficacy in various tumor models[34, 35].

In this study, we evaluated the utility of CBD-IL-12 therapy and the potency of a combination therapy involving CBD-IL-12 and Dox in mouse OS models. First, we show that CBD-IL-12 is selectively accumulated within the OS TME, due to the abundant expression of Collagen I and III in the OS. Next, we demonstrate that the combination therapy of CBD-IL-12 and Dox enhances therapeutic efficacy against multiple tumor models compared to monotherapy, which is mediated by the recruitment of DCs followed by maturation of cytotoxic T cells.

## Methods

### Mice and cancer cell lines

Female C57BL/6 mice (aged 6 to 8 weeks) were obtained from Charles River UK or Envigo and housed at the Hammersmith Central Biomedical service facility of Imperial College London. All the animals used at Imperial College London were handled in accordance with the 1986 Animal Scientific Procedures Act and under a United Kingdom Government Home Office-approved project license and overseen by ethical committees of Imperial College London. BVM03O cells were gifted from Prof. Brian A. Van Tine from Washington University. K7M2 cells (CRL-2836) were purchased from ATCC. BVM03O cells were maintained in IMDM medium (Gibco) supplemented with 20 % FBS (Gibco), 2 mM GlutaMAX (Gibco), 1% penicillin/streptomycin (Gibco), 1% MEM Non-essential amino acids (Gibco), and 3.5 μL of 2-mercaptoethanol. K7M2 cells were maintained in DMEM medium (Gibco) supplemented with 10 % FBS, 2 mM GlutaMAX, 1% penicillin/streptomycin.

### Production and purification of recombinant IL-12 and CBD-IL-12

Protein production was described previously[34]. In brief, to produce wild-type IL-12, optimized sequence encoding mouse p35 and p40 subunits were synthesized and subcloned into mammalian expression vector pcDNA3.1(+) by GenScript. 6xHistidine tag was added to the N terminus of the p35 subunit to enable affinity-based protein purification. To produce CBD-IL-12, a sequence encoding the CBD protein (A3 domain of VWF) was fused to the N terminus of the mouse p35 subunit to the C terminus of the mouse p40 subunit. 6xHistidine tag was added to the N terminus of the CBD-p35 subunit. Sequences encoding CBD-p35 and p40-CBD were subcloned into the mammalian expression vector pcDNA3.1(+) by GenScript. Recombinant IL-12 variants were produced by transient expression in HEK293F cells and purified as described previously. Purity of the proteins were evaluated using SDS-PAGE as described previously. Protein concentration was quantified by measuring absorbance at 280 nm using a NanoDrop One (Thermo Fisher Scientific). Full amino acid sequences of the IL-12 variants are available in the supplementary information.

### Data collection and Processing of the RNA-seq data

The RNA sequencing data were obtained from the GSE99671 dataset in the Gene Expression Omnibus (GEO) database (https://www.ncbi.nlm.nih.gov/geo/, accessed on 27^th^ May 2025). The raw gene expression counts were extracted for collagen type I alpha 1 chain (COL1A1) and collagen type III alpha 1 chain (COL3A1) genes. Expression values were log_2_-transformed using the formula log_2_(expression count + 1) to normalize the data distribution, which is standard practice for RNA-seq count data analysis. Expression data were visualized using box plots showing the distribution of log2-transformed expression values for each gene in normal versus tumor samples. Individual data points were overlaid on box plots to show the distribution of values across all samples.

### H & E staining and Masson’s trichrome staining

To prepare mouse tumor model, 3.0 x 10^6^ BVM03O cells were injected subcutaneously into the back skin of the female C57BL/6 mouse in 50 μL sterile PBS. After 50 days, the tumor was harvested and fixed with 2% PFA for 24 hours at 4 °C. The tumor pieces were rested in PBS for 24 hours at 4 °C, then following 24 hours incubation at 4 °C in the 70 % ethanol. Then the fixed tumors were sent to Research Histology Facility and perform H & E staining and Masson’s Trichrome Staining. In brief, after embedding in the paraffin, four-micrometers-thick mouse tissue sections were deparaffinized, rehydrated and H&E stained for the histological analysis. Collagen deposition was visualized using Masson’s Trichrome Stain (Diapath) following manufacturer’s instructions. The images of slides were acquired by slide scanner (Morphle Labs).

### Detection of collagen expression in mouse and human OS

To prepare mouse tumor model, 3.0 x 10^6^ BVM03O cells were injected subcutaneously into the back skin of the female C57BL/6 mouse in 50 μL sterile PBS. After 50 days, the tumor was harvested and fixed with 2% PFA for 24 hours at 4 °C. The tumor pieces were dehydrated by incubating with 30 % sucrose (in miliQ) for 24 hours to 48 hours. Once the tumor sunk, the tumor was embedded in the OCT compound (Agar Scientific) and frozen sections of 5 μm thickness were prepared. For the K7M2 tumor model, 5.0 x 10^6^ K7M2 cells were inoculated subcutaneously into the back skin of the female BALB/c mice in 50 μL sterile PBS. After 30 days, the tumor was harvested and prepare the frozen sections following same manner as BVM03O. Human OS cryosections were purchased from OriGene Technologies. For the collagen staining, cryosections were first blocked with 2% BSA in PBS at room temperature for 1 hour. Then, tissue samples were stained with rabbit anti-human/mouse collagen I antibodies (Abcam, ab21286) or rabbit anti-human/mouse collagen III antibodies (Abcam, ab184993) for 2 hours at room temperature. Tissue samples were then stained with the Alexa-647-labelled donkey anti-rabbit IgG (Jackson Immunoresearch Labs, 711-605-152) for 1 hour at room temperature. Sections were then covered with Prolong Gold Antifade Mountant containing 4,6-diamidino-2-phenylindole (DAPI; Thermo Fisher Scientific) and sealed with a coverslip. Microscopy was performed using a Leica SP8 confocal microscopy (Leica) at twenty times magnification, and images were processed using ImageJ software (NIH).

### Tumor accumulation in BVM03O-bearing mice

BVM03O tumor bearing mice received 416.6 pmol of CBD-IL-12 or 416.6 pmol of unmodified IL-12 (i.v., *n = 3*) when the tumor volume reached approximately 200 mm^3^. Tumors were harvested 30 minutes post treatment and were put into pre-weighed Lysing Matrix D tube (MP Biomedicals) containing 0.5 mL of T-PER Tissue Protein Extraction Reagent (Thermo Scientific) supplemented with cOmplete EDTA-free protease inhibitor cocktail (Roche). Tubes with tissue samples were weighed again and cut into small pieces using surgical scissors. The samples were lysed using FastPrep-24 5G (MP Biomedicals) and stored at -80°C until use. The amount of IL-12 was quantified using Mouse IL-12 p70 Uncoated ELISA Kit (Invitrogen). Unmodified IL-12 or CBD-IL-12 were used as standards. Total protein content was quantified using Pierce BCA Protein Assay kit (Thermo Fisher Scientific).

### Extraction of tumor-infiltrating lymphocytes

BVM03O cells (3 × 10^6^) were inoculated subcutaneously into the back skin of the female C57BL/6 mouse in 50 μl sterile PBS. Then, 35 days after tumor implantation, mice were treated with either PBS (i.v., *n = 5*). or CBD-IL-12 (i.v., *n = 4*). The tumor volume was measured once in 2 or 3 days. 13 days after the treatment, tumors were collected cut into small pieces using surgical scissors and digested for 30 min at 37 °C. Digestion medium was DMEM (Gibco) supplemented with 5% FBS, 2.0 mg/ml collagenase D (Sigma-Aldrich), 40 μg/ml DNase I (Roche). Single-cell suspensions were prepared using a 70 μm cell strainer (Thermo Fisher Scientific). Red blood cells were lysed with ACK lysing buffer (Gibco) for 5 minutes at room temperature and neutralized with DMEM 5% FBS.

### Analysis of cytotoxic T cells following CBD-IL-12 therapy in BVM03O tumor

The protocol for identifying immune cells was described previously[34]. Cells were stained with BD Horizon™ Fixable Viability Stain 510 (564406, BD Bioscience). Fc receptors were blocked with purified anti-mouse CD16/32 antibody (93, Biolegend). Cells were stained with a cocktail of anti-mouse antibodies and fixed with eBioscience IC Fixation Buffer (Invitrogen). Cells were acquired using a BD FACSymphony A3 flow cytometer and data were analyzed using FlowJo (BD). The following anti-mouse antibodies were used: CD45.2 APC-Cy7 (30-F11, Biolegend), CD3 BUV395 (145-2C11, BD), CD4 BUV805 (GK1.5, BD), CD8 AF700 (53-6.7, Biolegend), CD44 AF488 (IM7, Biolegend), CD62L PE Cy7 (MEL-14, Biolegend). Gating strategies are shown in Fig. S2.

### Anti-tumor efficacy of Dox against BVM03O tumor model mice

BVM03O tumor bearing mice were treated with PBS (i.v., *n = 7*), or 2 mg/kg of Dox (APExBIO) (i.v., *n = 7*) on day 35. Dox was additionally injected 3 and 6 days post initial treatment. 20 days after the initial treatment, the tumors were harvested for immunofluorescence staining. The tumor volume was calculated using the following formula: 4/3 x π x depth/2 × width/2 × hight/2.

### TUNEL assay

A classic TUNEL assay was performed with tumor frozen sections to detect and quantitate apoptotic cells using One-step TUNEL Apoptosis Assay Kit (Antibodies.com, A319767) according to manufacturer’s instructions. Three visual fields per TUNEL-stained slide were randomly selected for observation and photography under a confocal microscope (Leica). Image J software (NIH) was applied to calculate and analyze the density of apoptotic cells and average apoptotic index in each group.

### Immunofluorescence staining of Ki67 and CD31

Cryosections were first incubated with 0.3 % Triton X-100 for 15 minutes, then blocked with 2% BSA and 0.1 % Triton-X 100 in PBS at room temperature for 1 hour. Tissue samples were stained with rabbit anti- mouse Ki67 antibodies (Abcam, ab15580) and rat anti-mouse CD31 antibodies (BD, 553370) for overnight at 4 °C. After washing the slides with PBS-T 3 times, tissue samples were then stained with the Alexafluora-647-labelled donkey anti-rat IgG (Jackson Immunoresearch Labs, 712-606-153) and Alexafluora-594 labelled anti-rabbit IgG (Jackson Immunoresearch Labs, 711-585-152) for 1 hour at room temperature. After washing the slides with PBS-T 3 times, sections were then covered with Prolong Gold Antifade Mountant containing 4,6-diamidino-2-phenylindole (DAPI; Thermo Fisher Scientific) and sealed with a coverslip. Microscopy was performed using a Leica SP8 confocal microscopy (Leica).

### Combination therapy of CBD-IL-12 and Dox against OS model

BVM03O tumor bearing mice were treated with PBS (i.v., *n* = 5), 416.6 pmol of CBD-IL-12 (i.v., *n* = 5), and 416.6 pmol of CBD-IL-12 + 2 mg/kg of Dox (i.v., *n* = 5) on day 35 post tumor inoculation. Dox was additionally injected on 3 and 6 days post initial treatment. K7M2 tumor bearing mice were treated with PBS (i.v., *n* = 4) and 416.6 pmol of CBD-IL-12 + 2 mg/kg of Dox (i.v., *n* = 3) once the tumor volume reaches 50 mm^3^. Dox was additionally injected on day 3 and day 6. The tumor volume was measured using a digital caliper and the volume was calculated using the following formula: 4/3 x π x depth/2 × width/2 × hight/2). Mice were euthanized when tumor volume had exceeded 600 mm^3^ or tumor ulceration of more than 7 mm in diameter had been observed.

### Analysis of tumor-infiltrating lymphocytes in combination therapy with CBD-IL-12 and Dox

BVM03O tumor bearing mice were treated with PBS (i.v., *n* = 5), 416.6 pmol of CBD-IL-12 (i.v., *n* = 5), and 416.6 pmol of CBD-IL-12 + 2 mg/kg of Dox (i.v., *n* = 5) on day 35. Dox was additionally injected 3 and 6 days post initial treatment, then the tumors were harvested 13 days post initial treatment.

The protocol for identifying immune cells was described previously[34]. After extracting tumor-infiltrating lymphocytes, Cells were stained with LIVE/DEAD™ Fixable Near IR (876) Viability Kit, for 808 nm excitation (Thermofisher). Fc receptors were blocked with purified anti-mouse CD16/32 antibody (93, Biolegend). Cells were stained with a cocktail of anti-mouse antibodies and fixed with eBioscience IC Fixation Buffer (Invitrogen). The cells were washed with eBioscience™ Permeabilization Buffer, then the intracellular staining was performed. The cells were acquired using CytoFLEX LX flow cytometer (Beckman) and the data were analyzed using FlowJo (BD). The following anti-mouse antibodies were used for T cells: CD45.2 APC-Cy7 (30-F11, Biolegend), CD3 BUV395 (145-2C11, BD), CD4 AF700 (GK1.5, Biolegend), CD8 PE Duzzle 594 (53-6.7, Biolegend), KLRG1 BV510 (2F1/KLRG1, Biolegend), CD127 PE-Cy5 (A7R34, Biolegend), CD11b APC (M1/70, Biolegend), Ly6G BVV737 (1A8, BD), Ly6C AF488 (HK1.4, Biolegend), F4/80 PE-Cy7 (BM8, Biolegend), MHCII BUV805 (M5/114.15.2), CD11c BV421 (N418, Biolegend), XCR1 BV650 (ZET, Biolegend), SIRPα PE (P84, Biolegend), CD38 BV786 (90/CD38, BD), and CD206 BV605 (C068C2, Biolegend). Gating strategies are shown in Fig. S3 and S4.

### Statistical analysis

Statistical analysis was performed using GraphPad Prism 10. Tests used were indicated in figure captions.

## Results

### Collagen type I and collagen type III are abundant in OS tissue

To investigate collagen expression patterns in OS, we first analyzed RNA sequencing data from the Gene Expression Omnibus (GEO) database (accession GSE99671), which contains transcriptomic profiles of human OS samples (n=18) and matched non-tumoral bone tissue (n=18). The mean expression of *COL1A* (coding for chain alpha 1 of collagen I) and *COL3A1* (coding for chain alpha 1 of collagen III) was visualized in a box plot (Fig. 1a, b). While *COL1A1* expression showed no statistical difference compared to normal bone tissue (Fig. 1a), the expression of *COL3A1* was significantly upregulated in the OS samples (Fig. 1b).

**Figure. 1.**
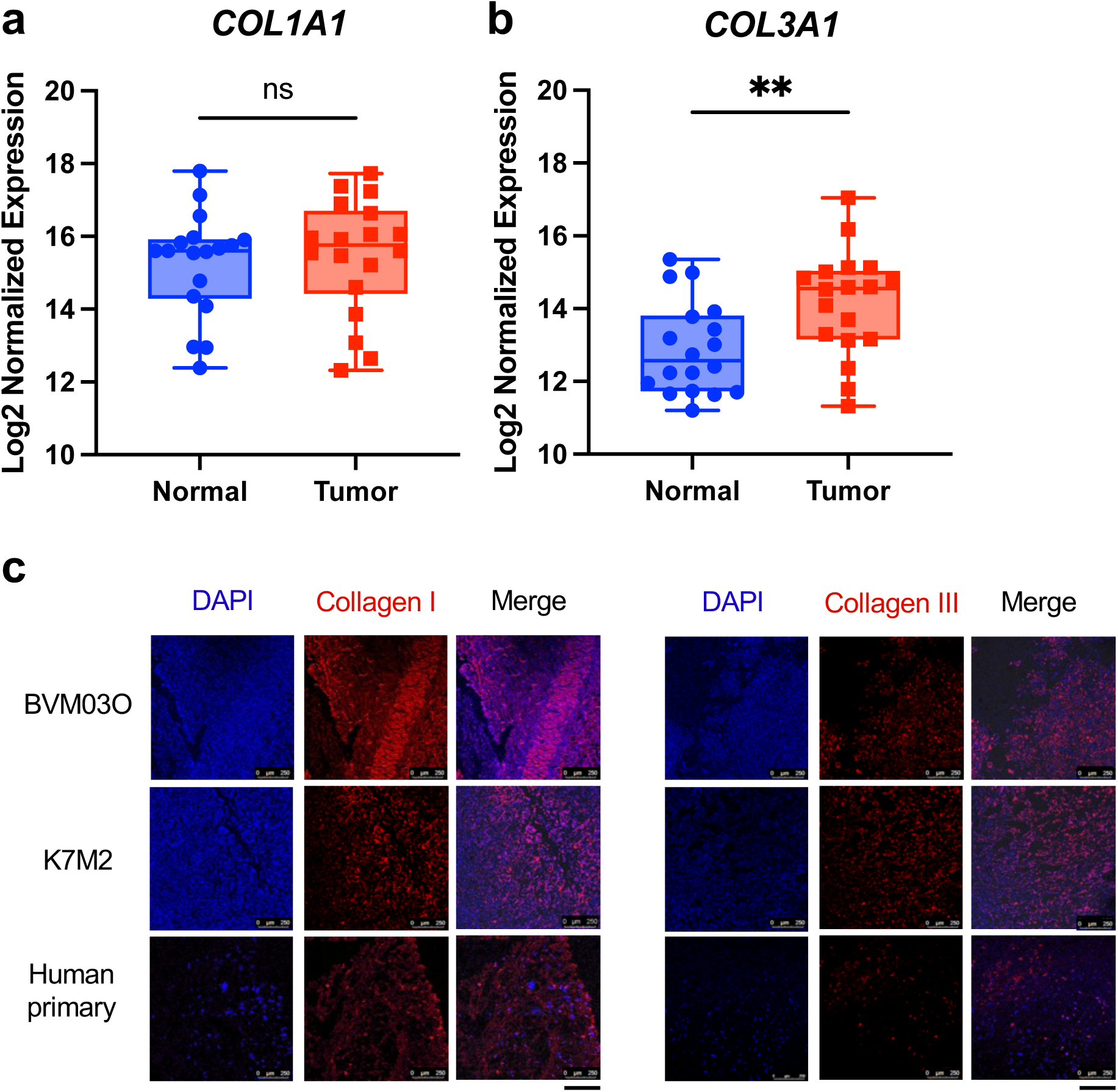
CBD can target OS tissue. The box plot of (a) collagen type I and (b) collagen type III expression in OS tissue compared with paired normal bone samples. (c) Collagen type I and III expression in both mouse (BVM03O and K7M2) and human OS. The cryosections were imaged by confocal microscopy. Nucleus and collagen type I or collagen type III were blue and red, respectively. Scale bars, 250 μm. The results are expressed as mean ± S.E.M. Statistical analysis was performed using an unpaired t-test for (a) and (b). *P < 0.05, **P < 0.01. N.S., not significant; PBS, phosphate-buffered saline.

To also confirm collagen expression in mouse pre-clinical models, we established subcutaneous OS models using two complementary murine cell lines namely BVM03O and K7M2 with distinct biological characteristics. BVM03O represents a primary OS model derived from mouse osteoblast harboring p53 mutation[36], recapitulating the genetic alterations commonly observed in human OS[37]. K7M2, is originally derived from OS that metastasized to the lungs, providing a model of advanced disease[38]. Previous studies have demonstrated that K7M2 tumors exhibit partial responsiveness to checkpoint inhibitor therapy without achieving complete remission, mirroring the limited efficacy of current immunotherapeutic approaches in clinical trials[39]. Histopathological evaluation of the BVM03O tumors via HE stains revealed diverse morphologies among tumour cells. Notably, the presence of homogeneous glassy material reminiscent of osteoid formation in bone tissues was evident (Fig. S1a). Additionally, trichrome staining showcased an extensive blue hue throughout the area, indicating the potential presence of intratumoural collagen within the tumor (Fig. S1b).

Immunofluorescence analysis of subcutaneous K7M2 tumors confirmed abundant expression of both collagen type I and type III throughout the tumor microenvironment in both murine models (Fig. 1c). Similar collagen deposition patterns were observed in human OS tissue samples, with extensive collagen networks distributed throughout the tumor stroma (Fig. 1c), validating our transcriptomic findings. The widespread expression of both collagen subtypes across primary and metastatic OS models rationalizes collagen-targeted drug delivery as a therapeutic strategy for OS treatment.

### CBD fusion enhances the accumulation of IL-12 within OS TME and induces intratumoral inflammation

We validated collagen-targeted therapy of CBD-IL-12 against OS. We firstly injected CBD-IL-12 or unmodified IL-12 to BVM03O-bearing mice intravenously, and tumors were harvested at 30 minutes post treatment. As hypothesized, CBD-IL-12 showed 4 times higher tumor accumulation than unmodified IL-12, suggesting that CBD fusion enhanced the tumor targeting of IL-12 in the OS model (Fig. 2a). Then, we tested therapeutic effect of CBD-IL-12 using BVM03O OS model (Fig. 2b). Although not significant, the intravenous administration of CBD-IL-12 had a trend to suppress the tumor growth until 8 days post treatment, after which the tumor continued to grow gradually until day 13. We next investigated immune infiltrates in the tumor. The frequency of CD8^+^ T cells within CD45^+^ cells was 0.7% in the PBS-treated group. CBD-IL-12 therapy demonstrated a 2.2-fold increase in the CD8^+^ T cells within CD45^+^ cells, suggesting that CBD-IL-12 enhanced immune cells recruitment in the TME (Fig. 2c). Interestingly, the percentage of effector-memory CD8^+^ T cells exhibited a 2-fold increase in the CBD-IL-12-treated group (Fig. 2d), which is consistent with previous study using different types of tumor models[34]. CBD-IL-12 treatment did not change the number of CD4^+^ T cells, which is consistent with a previous report (Fig. S2a)[34]. This suggests that CBD-IL-12 therapy induced inflammation in an immunologically cold OS tumor.

**Figure. 2.**
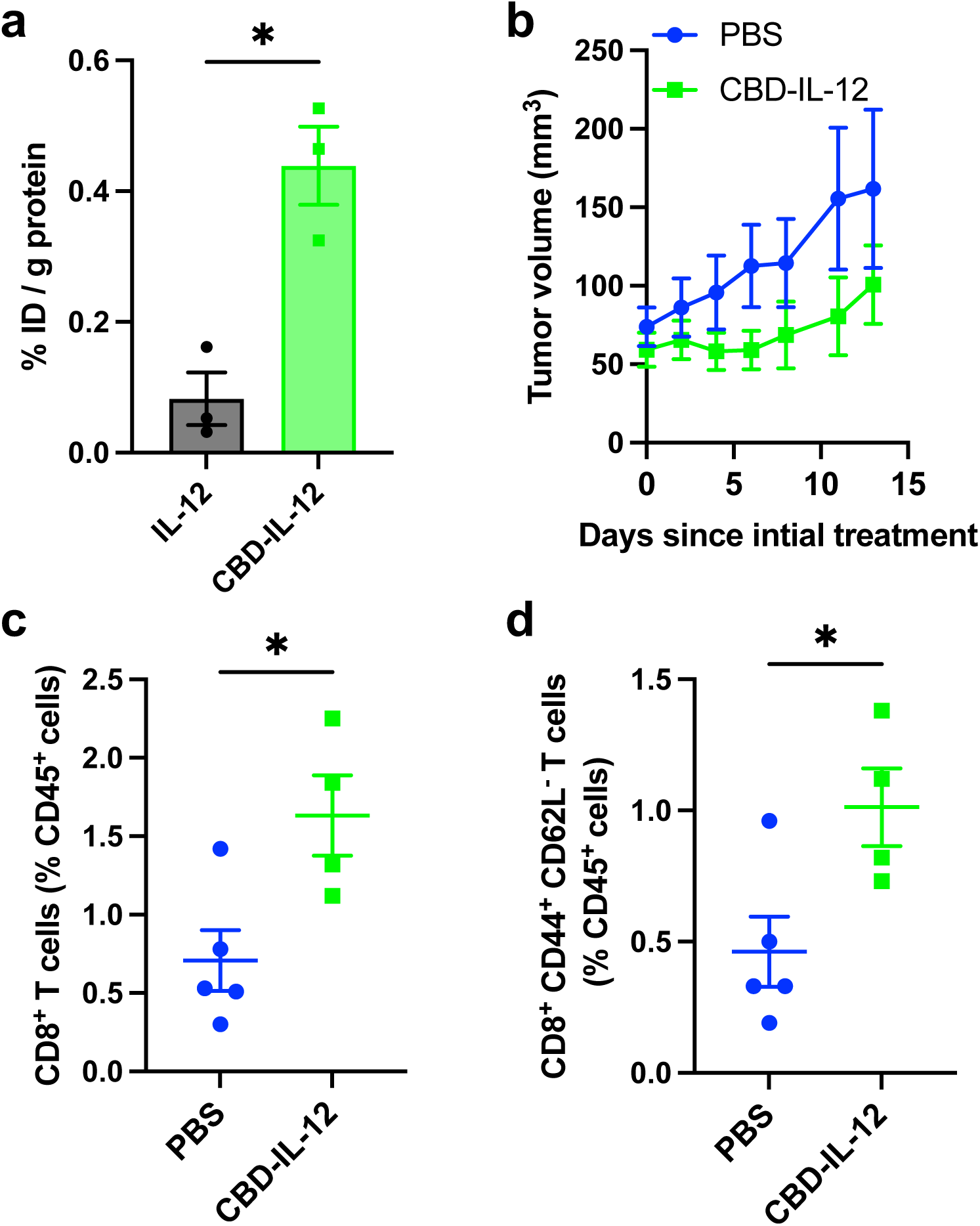
CBD-IL-12 treatment increases cytotoxic CD8^+^ T cells in the BVM03O OS. (a) BVM03O cells (3 × 10^6^) were inoculated subcutaneously. When the tumor volume reached 200 mm^3^, 416.6 pmol of CBD–IL-12 or 416.6 pmol of unmodified IL-12 was injected intravenously. After 30 minutes, the tumor was harvested, and the amount of IL-12 was quantified by ELISA (*n* = 3). (b) We inoculated mice with 3 × 10^6^ BVM03O OS cells subcutaneously on the back skin, and the mice were treated with PBS (i.v., *n* = 5), 416.6 pmol of CBD-IL-12 (i.v., *n* = 4) on day 35. Tumors were collected on day 48, followed by flow cytometric analysis. The average tumor growth curve is shown in (a). Frequency of (c) CD3^+^CD8^+^ and (d) CTL (CD3^−^CD8^+^CD44^+^CD62L^-^) tumor-infiltrating T cells within CD45^+^ leukocytes. The results are expressed as mean ± S.E.M. Statistical analysis was performed using an unpaired t-test for (c) and (d). *P < 0.05, **P < 0.01. N.S., not significant; PBS, phosphate-buffered saline.

### Dox did not suppress the tumor growth, but induced apoptosis in the OS tumor

To confirm the efficacy of Dox in our OS model, we treated the BVM03O bearing mice with either PBS (i.v. *n = 7*) or Dox (i.v. *n = 7*) once every 3 days for three times. Dox monotherapy had little effect on the tumor growth (Fig. 3a), which is consistent with clinical setting[40]. To further assess the efficacy of Dox therapy, the tumors were harvested to evaluate apoptosis in the tumors 20 days after the initial Dox treatment. After fixation of the tumors, we performed Terminal deoxynucleotidyl transferase dUTP nick-end labelling (TUNEL) assay to evaluate the apoptosis (Fig. 3b). TUNEL assay revealed that Dox therapy showed 3 times higher frequency of merged area of nucleus and TUNEL than the PBS-treated group (Fig. 3c), indicating that Dox induced apoptosis in the tumor.

**Figure. 3.**
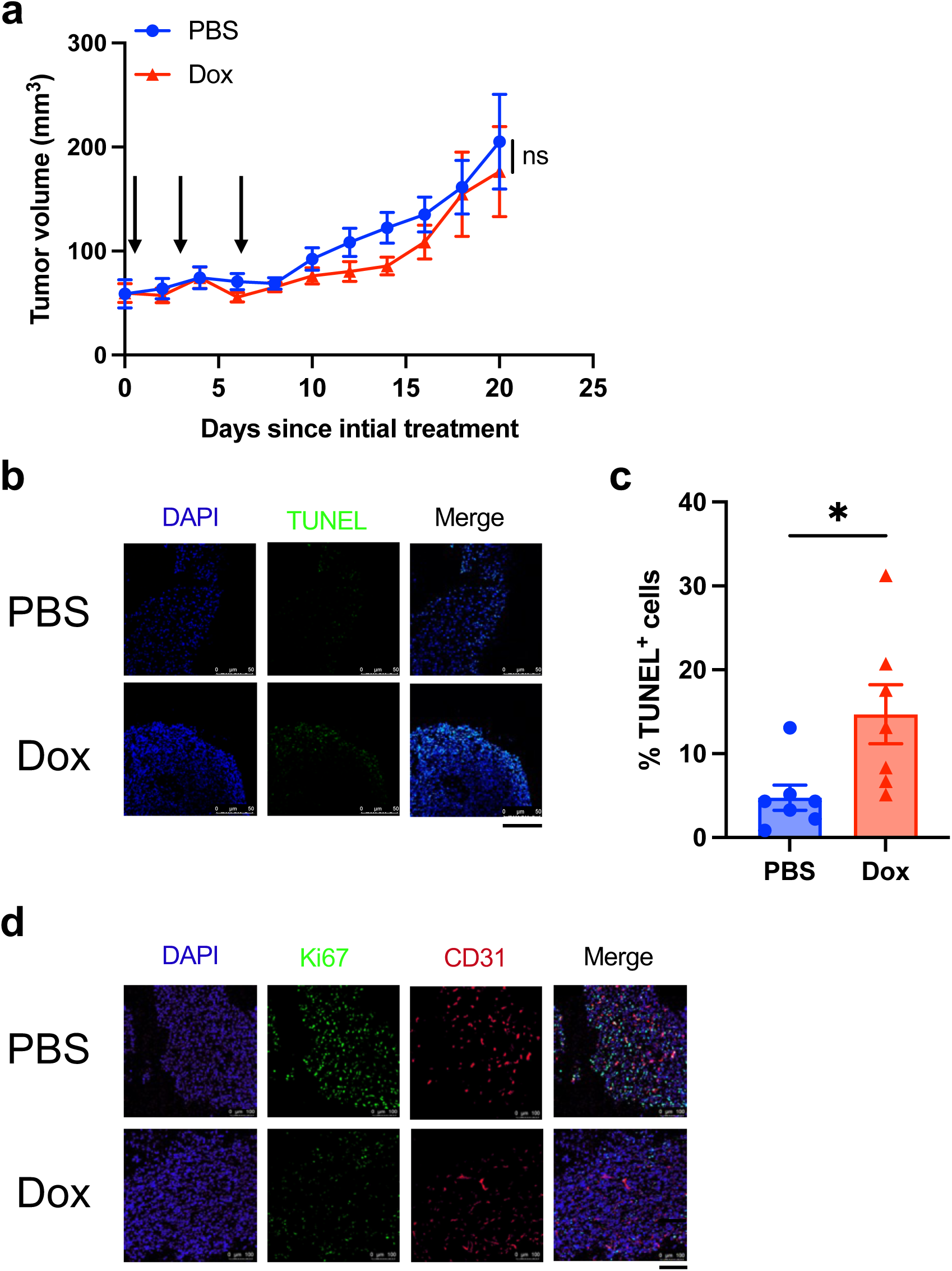
Dox treatment can induce cytotoxic apoptosis in BVM03O OS. We inoculated mice with 3 × 10^6^ BVM03O OS cells subcutaneously on the back skin and the mice were treated with PBS (i.v., *n* = 7), 2 mg/kg of Dox (i.v., *n* = 7) on day 35. Dox was additionally injected 3 and 6 days 6 post initial treatment. On day 20 after initial treatment, the tumor was harvested for immunofluorescence staining. The average tumor growth curve was shown in (a). (b) TUNEL staining was conducted to study tumor apoptosis. Scale bars, 50 μm. (c) The average percentage of TUNEL-positive cells. (d) Ki67 staining was performed in tumor tissue sections of different groups to visualize tumor proliferation. CD31 staining was applied to analyze tumor angiogenesis. Scale bars, 100 μm. The results are expressed as mean ± S.E.M. Statistical analysis were performed using unpaired t test for (a) and unpaired t test with Welch’s correction for (d). *P < 0.05, N.S., not significant; PBS, phosphate-buffered saline.

Dox therapy has been demonstrated to suppress Ki67 and CD31 expression, which serve as markers of proliferation and angiogenesis, respectively[41]. We analyzed the Ki67 and CD31 expression through immunofluorescence staining (Fig. 3d). As reported, Dox therapy decreased the expression of both Ki67 and CD31, confirming that dox has an effect on the TME at the molecular level. Since the BVM03O cells do not express p53[36], a lower dose of Dox might be sufficient to induce apoptosis and reduce proliferation and angiogenesis, but not to demonstrate significant therapeutic efficacy and induce OS tumor growth inhibition[42]. This is likely because the absence of p53 compromises DNA damage–induced cell-cycle arrest and sustained apoptotic signaling, which are required for durable tumor regression in vivo[43].

### CBD-IL-12 showed a synergistic effect with Dox

Despite our demonstration that Dox monotherapy induced apoptosis (Fig. 3b, c), this treatment did not suppress tumor growth in OS tumour models, which is representative of its insufficient clinical efficiency and highlights a need for combination strategy (Fig. 3a). To achieve superior therapeutic efficacy, we evaluated the synergistic effect of CBD-IL-12 and Dox combo therapy in OS models. Thirty-five days post inoculation with BVM03O cells, the mice were treated with PBS (i.v., *n* = 5), CBD-IL-12 (i.v., *n* = 5), or CBD-IL-12 + Dox (i.v., *n* = 5) (Fig. 4a, b). CBD-IL-12 therapy initially suppressed tumor growth, but the tumor started to regrow 1 week post treatment. In contrast, CBD-IL-12 in combination with Dox therapy suppressed tumor growth until day 56, which is an improvement in the medial survival time of 12 days and 6 days compared to PBS and CBD-IL-12, respectively (Fig. 4c). These data suggested that CBD-IL-12 therapy exhibited a synergistic effect with Dox therapy. We then assessed the generalizability of our findings by testing the combination therapy in a second OS model based on the syngeneic OS cell line K7M2. These cells were inoculated into the back skin of the mice and treated with either PBS (i.v., *n* = 4) or CBD-IL-12 + Dox (i.v., *n* = 3) once the average tumor volume reached 50 mm^3^. The combination therapy suppressed tumor growth for 2 weeks post treatment (Fig. 4d, e). Consequently, the therapy extended median survival 7 days compared to PBS-treated group (Fig. 4f). Together, these results demonstrated that the combination of CBD-IL-12 with Dox exhibits therapeutic efficacy against two different mouse tumor models.

**Figure. 4.**
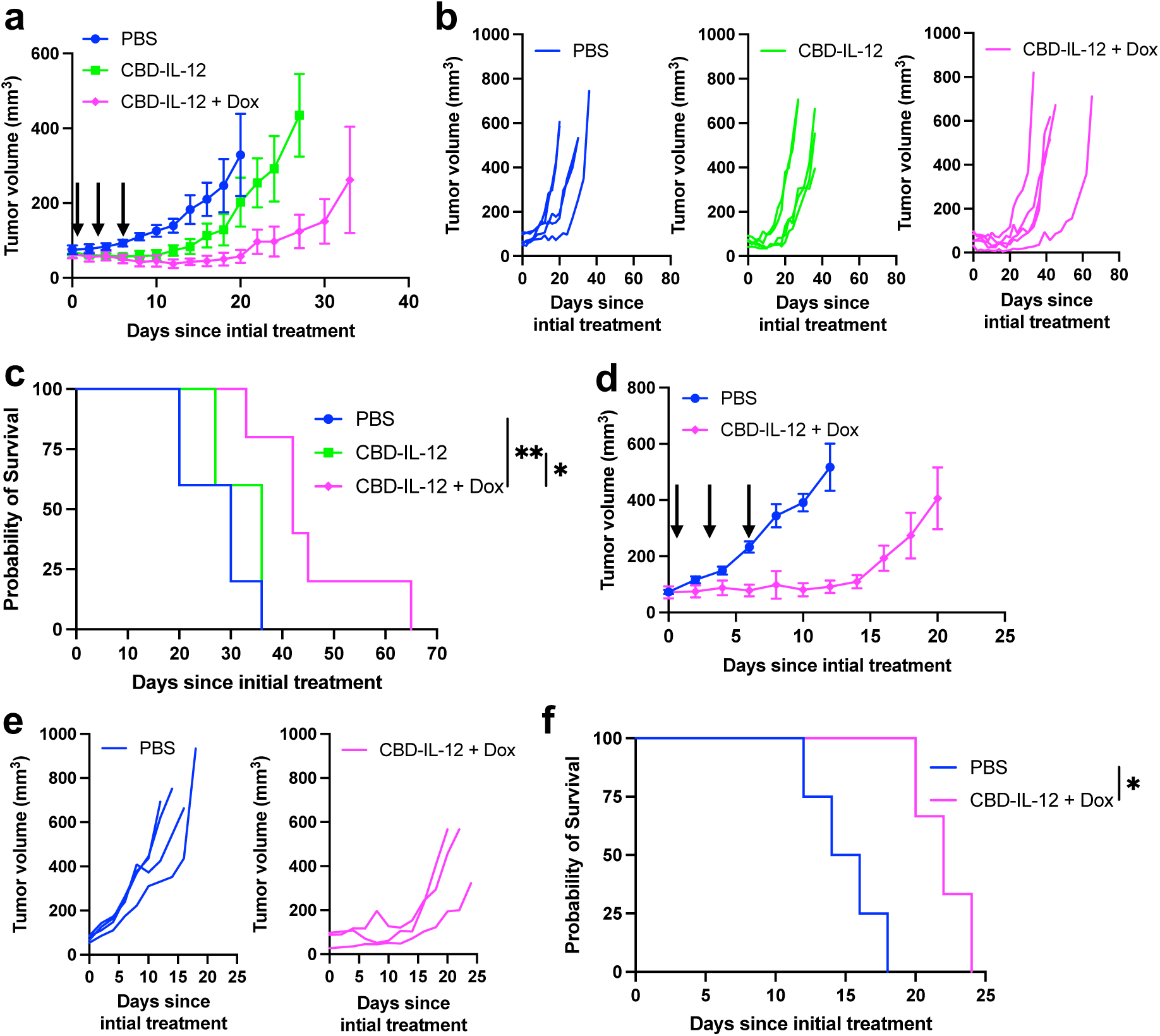
CBD-IL-12 synergizes with Dox and regresses OS. We inoculated mice with 3 × 10^6^ BVM03O OS cells subcutaneously on the back skin and the mice were treated with PBS (i.v., *n* = 5), 416.6 pmol of CBD–IL-12 (i.v., *n* = 5), and 416.6 pmol of CBD–IL-12 + 2 mg/kg of Dox (i.v., *n* = 5) on day 35. Dox was additionally injected 3 and 6 days post initial treatment. Average tumor growth curve (a), individual tumor curves (b), and survival curves (c) are shown. We inoculated mice with 5 × 10^6^ K7M2 OS cells subcutaneously on the back skin and the mice were treated with PBS (i.v., *n* = 4) and 416.6 pmol of CBD-IL-12 + 2 mg/kg of Dox (i.v., *n* = 3) once the tumor volume reaches 50 mm^3^. Dox was additionally injected on day 3 and day 6. Average tumor curve (d), individual tumor curves (e), and survival curves (f) are shown. The results are expressed as mean ± S.E.M. Log-rank test was used for (c) and (f) in the statistical analysis. *P < 0.05, **P < 0.01. N.S., not significant; PBS, phosphate-buffered saline.

### CBD-IL-12 + Dox combination therapy enhanced anti-tumor immunity compared to Dox solo therapy

To uncover the mechanism behind the therapeutic efficacy of CBD-IL-12 + Dox combination therapy, we firstly characterized T cells population in BVM03O-bearing mice. We injected Dox (i.v., *n* = 5) or CBD-IL-12 + Dox (i.v., *n* = 5) intravenously and extracted lymphocytes from the tumor 13 days post treatment. Although not significant, CBD-IL-12 + Dox combination therapy suppressed tumor growth (Fig. 5a). CBD-IL-12 + Dox increased total CD45^+^ cells within the tumor compared to Dox solo therapy (Fig. 5b), whereas CD4^+^ and CD8^+^ T cells were maintained in both groups (Fig. 5c and d). Interestingly, the number of killer cell lectin like receptor G1 (KLRG1)^+^ CD127^-^ CD8^+^ T cells which have a potent immediate killing capacity[44], were significantly increased in the combination therapy group (Fig. 5e). CBD-IL-12 + Dox combo therapy increased total number of CD11c^+^ Major Histocompatibility Complex Class II (MHCII)^+^ conventional DCs (cDCs) compared to Dox monotherapy (Fig. 5f). Furthermore, the combo therapy enhanced X-C Motif Chemokine Receptor 1 (XCR1)^+^ cDC1 in the tumor (Fig. 5g). cDC1 have been implicated in activating anti-tumor cytotoxic CD8^+^ T cells, which might have been recruited by CBD-IL-12[45]. Moreover, CBD-IL-12 + Dox therapy exhibited higher M1/M2 ratio compared to Dox therapy (Fig. 5h), which is one of the predictors of therapeutic efficacy[46, 47]. These data suggest that combining CBD-IL-12 therapy with Dox monotherapy increases cDC1. This ultimately promotes the of effector T cells, leading to the tumor growth suppression.

**Fig. 5.**
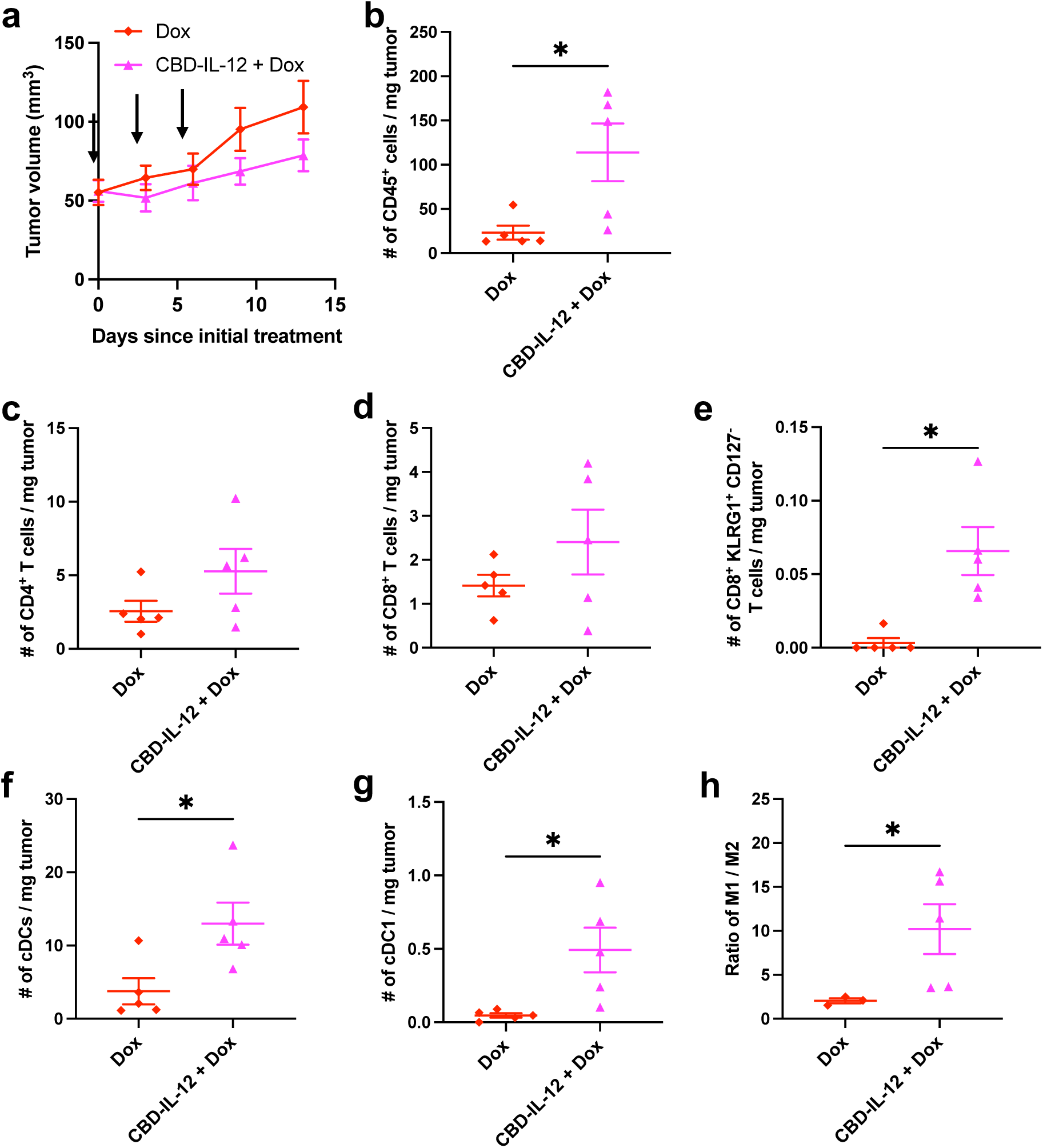
Dox + CBD-IL-12 combination therapy increase cytotoxic CD8^+^ T cells and DCs in the BVM03O OS. (a) BVM03O cells (3 × 10^6^) were inoculated subcutaneously. When the tumour volume reached 50 mm^3^, 2 mg/kg of Dox (i.v., *n* = 5) or 2 mg/kg of Dox and 416.6 pmol of CBD–IL-12 (i.v., *n* = 5) were injected intravenously on day 36. Dox was additionally injected 3 and 6 days post initial treatment. Tumors were collected 13 days post initial treatment, followed by flow cytometric analysis. The average tumour growth curve was shown in (a). Counts of (b) total CD45^+^, (c) CD3^+^CD4^+^, (d) CD3^+^CD8^+^, (e) CD3^+^CD8^+^KLRG1^+^CD127^-^, (f) total cDCs, (g) cDC1 were shown. The ratio of M1-like macrophage (F480^+^CD38^+^CD206^-^) versus M2-like macrophage (F480^+^CD38^-^CD206^+^) was shown for subjects with detectable macrophage populations in (h). Two mice in the Dox were excluded from ratio analysis as they exhibited zero detectable M1-like and M2-like macrophages. The results are expressed as mean ± S.E.M. Statistical analysis were performed using unpaired t test for (f), and unpaired t test with Welch’s correction was used for (b), (e), (g), (h).

## Discussion

A promising strategy in the treatment of different types of cancer is chemoimmunotherapy, which combines chemotherapy and immunotherapy[48]. Chemotherapy exerts its anti-tumor effect by killing cancer cells via ICDs, which is a process that is facilitated by the release of tumor antigens and then stimulate immune responses[48, 49]. Immunotherapy serves to enhance the anti-tumor effect initiated by chemotherapy, thereby potentiating its therapeutic efficacy[49]. Despite their clinical success, therapeutic efficacy of the chemoimmunotherapy is inconsistent across different cancer types[50]. T cell dysfunction and the immunosuppressive phenotype of the tumor represent significant challenges, as functional immune cells are necessary for tumor cells to be killed in the chemoimmunotherapy[49]. To address these concerns, IL-12, which activates and recruits various immune cells, is a candidate when administrated in conjunction with chemotherapy regimens, including Dox. Although IL-12 and Dox chemoimmunotherapy have been shown to be effective in cancer models, current therapeutic approaches are inadequate in demonstrating sufficient therapeutic efficacy with minimal immune adverse effects, due to the absence of targeted strategies[30]. We demonstrated that CBD-IL-12 + Dox chemoimmunotherapy would be among the first examples of collagen-targeted delivery enhancing anti-tumor effect and synergizing with Dox against Dox-unresponsive OS models.

Our characterization of collagen expression in OS provides molecular basis of this targeted approach. We demonstrated an abundant collagen I and III in OS TME, which has been recognized as one of the therapeutic difficulties[51], as this hampered immune infiltration in the TME[52]. The 4-fold enhanced tumor accumulation of CBD-IL-12 compared to unmodified IL-12 suggests that collagen-rich microenvironment of OS would become a retention substrate of CBD-IL-12[53, 54]. Simultaneously, CBD-IL-12 therapy overcome the immunosuppressive microenvironment of OS, which was confirmed by the achievement of 2.2-fold increase in the CD8^+^ T cells and doubled effector-memory CD8^+^ T cells in the TME. Despite of the improvement of immune microenvironment, CBD-IL-12 monotherapy demonstrated transient tumor growth suppression with regrowth by day 8. This might be due to the lower baseline of CD8^+^ T cells in the OS compared to other cancer models[34]. These results imply both the utility of CBD-IL-12 therapy, as well as the need for combination strategies in the treatment of OS.

Although Dox monotherapy does not suppress the tumor growth of OS, combination therapy utilizing both CBD-IL-12 and Dox significantly extended the median survival of BVM03O and K7M2 OS models. Mechanistic analysis has revealed that the number of KLRG1^+^ CD127^-^ CD8^+^ T cells and cDC1 are increased in combo therapy. This suggests that CBD-IL-12 therapy induced the recruitment of cDC1, which might have cross-presented tumor antigens to the CD8^+^ T cells, thereby contributing for maturation. Furthermore, the elevated of M1/M2 ratio within the TME signifies that M1-like macrophages may engulf apoptotic cells which have increased as a result of Dox therapy. This comprehensive immune remodeling distinguishes the combination therapy from monotherapies and validates the complementary nature of ICD induction with sustained IL-12-mediated immune activation.

In comparison with previous preclinical studies that utilized combination of Dox and IL-12 as a chemoimmunotherapy approach[55, 56], our collagen-targeted strategy enhances the localization of IL-12, achieving sufficient therapeutic efficacy with minimum dose of IL-12. IL-12 recombinant protein and Dox-loaded nanoparticle demonstrated the therapeutic efficacy in the hepatocellular carcinoma model with M1 polarization *in vitro*[56]. Our data demonstrate not only an augmented M1/M2 ratio, but also more cytotoxic T cells and antigen presenting cDC1 cells compared with Dox monotherapy in the OS preclinical model. This finding underscores the mechanism underlying the combination therapy of CBD-IL-12 and Dox, thereby providing a novel chemoimmunotherapy approach in the OS.

## Conclusion

The combination therapy of CBD-IL-12 and Dox represents a mechanistically rational approach against OS. Utilizing CBD-mediated strategy, CBD-IL-12 exhibits preferential tumor accumulation and activates the immunologically “cold” microenvironment in OS TME. Combination therapy with Dox further enhanced therapeutic efficacy, which potentially emerges as a promising clinical candidate warranting translational investigation for next-generation OS treatment protocols.

## Acknowledgements

We thank the LMS/NIHR Imperial Biomedical Research Centre Flow Cytometry Facility for its support. We thank Research Histology Facility at National Heart and Lung Institute for its support. This research was funded in part by Sarcoma UK (SUK23.2024 to J.I.), Bone Cancer Research Trust (BCRT/9824 to J.I.), Pancreatic Cancer Research Fund (PCRF 2022 (A010339) to J.I.) and The Brain Tumour Charity (QfC_2022_10657 to J.I.).

## CRediT authorship contribution statement

**Takumi Matsuo**: Writing – original draft, Visualization, Validation, Methodology, Investigation, Formal analysis, Data curation, Conceptualization. **Laurine Noblecourt**: Writing – review & editing, Methodology, Investigation. **Pooja Kaur**: Writing – review & editing, Methodology, Investigation. **Chuyi Wang**: Methodology, Investigation. **Po-Chuan Chiu**: Methodology, Investigation. **Koichi Sasaki**: Writing – review & editing, Methodology, Investigation. **Charanjit Singh**: Methodology. **Adrian Larkeryd**: Methodology. **Anguraj Sadanandam**: Methodology. **Paul H. Huang**: Writing – review & editing, Supervision, Resources, Project administration, Methodology, Investigation, Funding acquisition, Conceptualization. **Jun Ishihara**: Writing – review & editing, Validation, Supervision, Resources, Project administration, Methodology, Investigation, Funding acquisition, Formal analysis, Data curation, Conceptualization.

## Additional Information

Requests for materials should be addressed to Jun Ishihara.

---

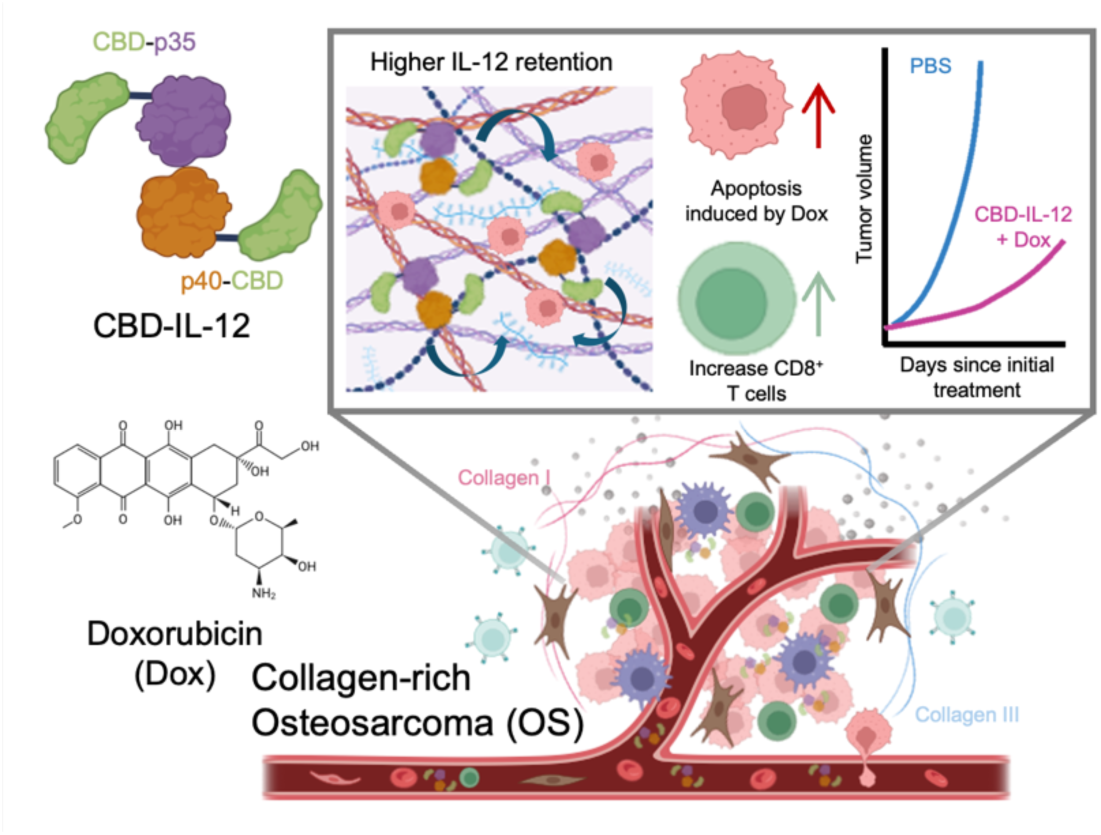
For graphical abstract only.

