## Supplementary Figures for "Collagen targeting IL-12 combined with Doxorubicin enhances the anti-tumor effect against osteosarcoma"

Supplementary Figure. S1. Representative images of H & E staining and Masson’s Trichrome staining of BVM03O tumor.

Supplementary Figure. S2. The frequency of $CD4^+$ T cells in $CD45^+$ leukocytes.

Supplementary Figure. S3. Gating strategy for characterizing T cells.

Supplementary Figure. S4. Gating strategy for characterizing T cells in the CBD-IL-12 + Dox combo therapy.

Supplementary Figure. S5. Gating strategy for characterizing myeloid cells in the CBD-IL-12 + Dox combo therapy.

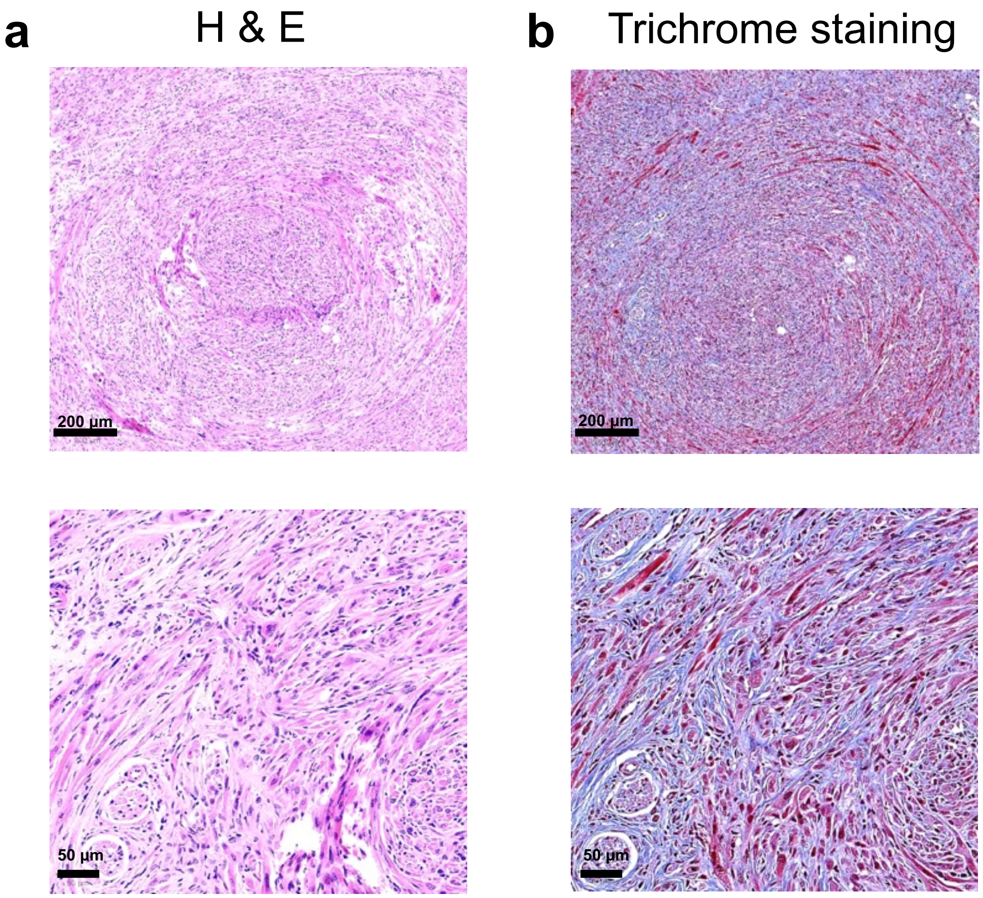


**Figure S1**. (a) Representative histological analysis of BVM03O tumor lesions in Hematoxylin & eosin. (b) Representative images of Masson's Trichrome staining of BVM03O tumor.

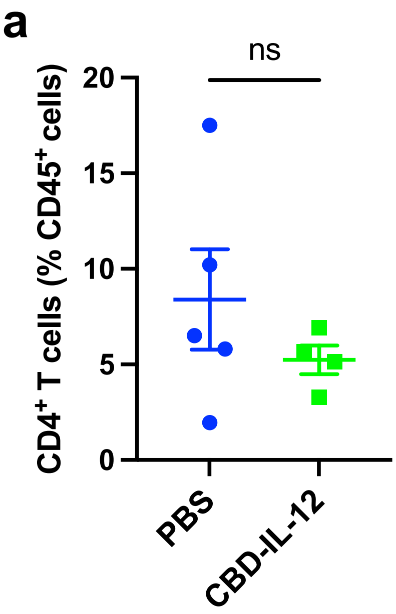


**Figure S2**. We inoculated mice with 3 × $10^6$ BVM03O OS cells subcutaneously on the back skin, and the mice were treated with PBS (i.v., *n* = 5), 416.6 pmol of CBD-IL-12 (i.v., *n* = 4) on day 35. Tumors were collected on day 48, followed by flow cytometric analysis. (a) $CD3^+CD4^+$ tumor-infiltrating T cells within $CD45^+$ leukocytes.

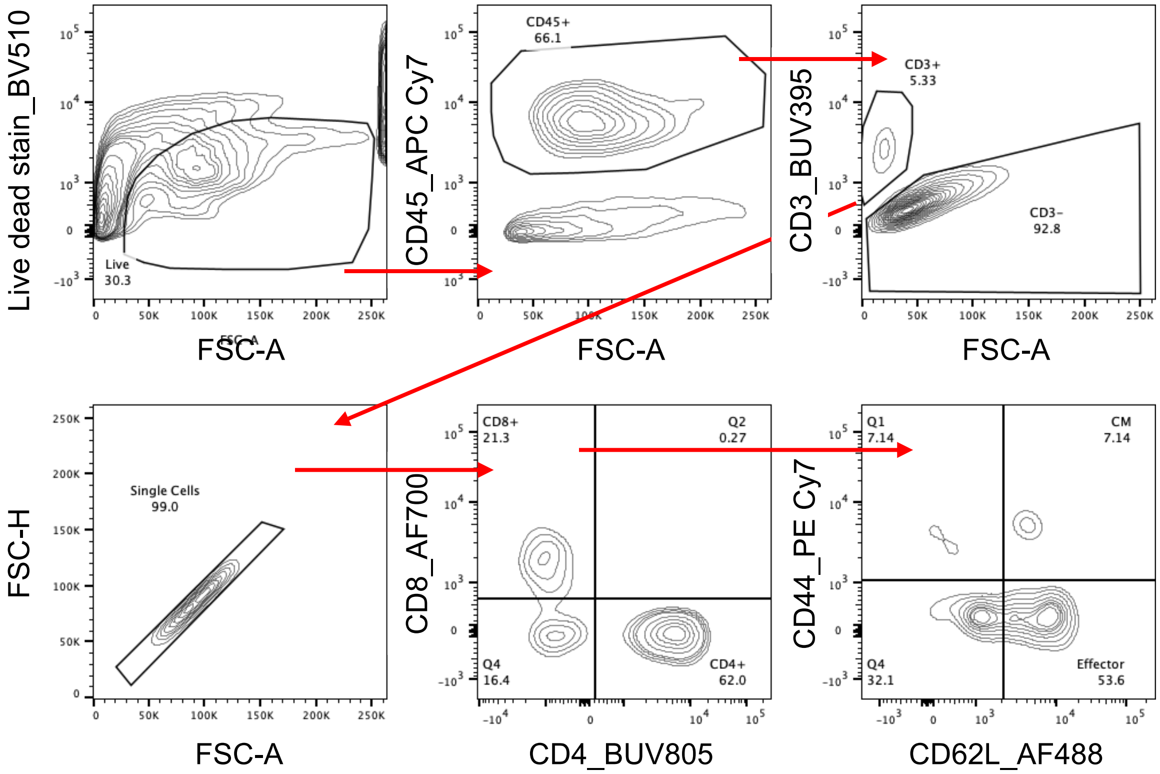


**Figure S3.** Gating strategy for characterizing T cells. Cells were first gated based on BD Horizon Fixable Viability Stain 510 followed by staining with indicated markers.

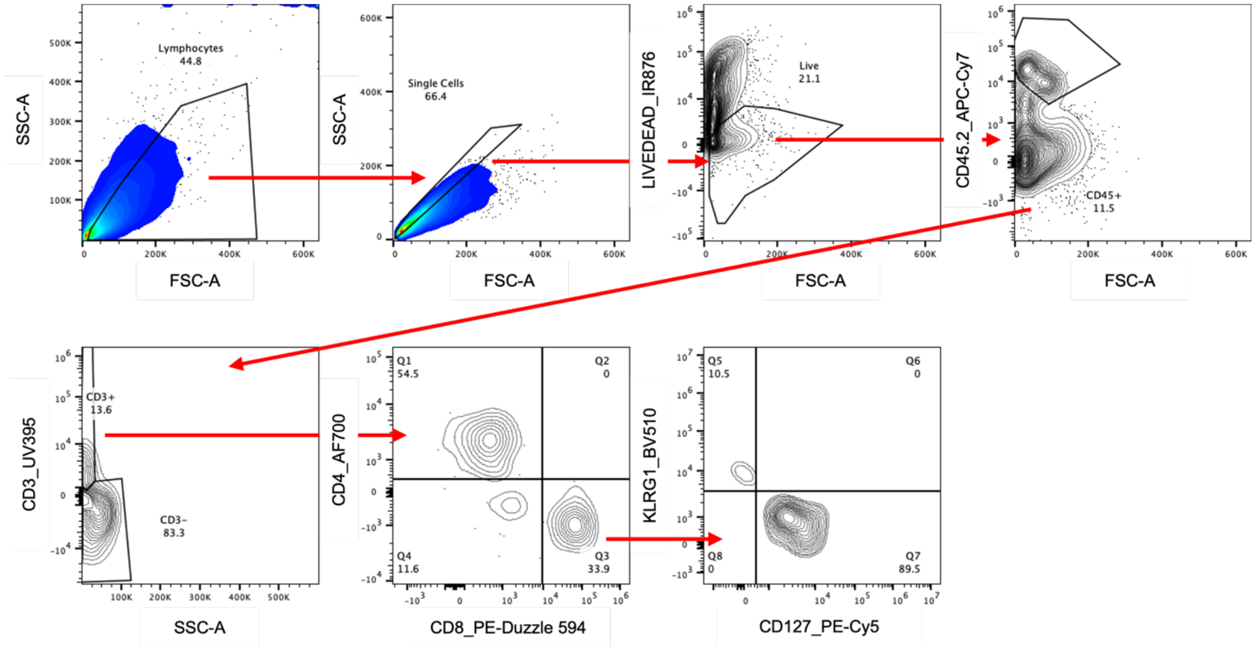


**Figure S4.** Gating strategy for characterizing T cells in the CBD-IL-12 + Dox combo therapy. Cells were first gated based on LIVE/DEAD™ Fixable Near IR (876) Viability Kit followed by staining with indicated markers.

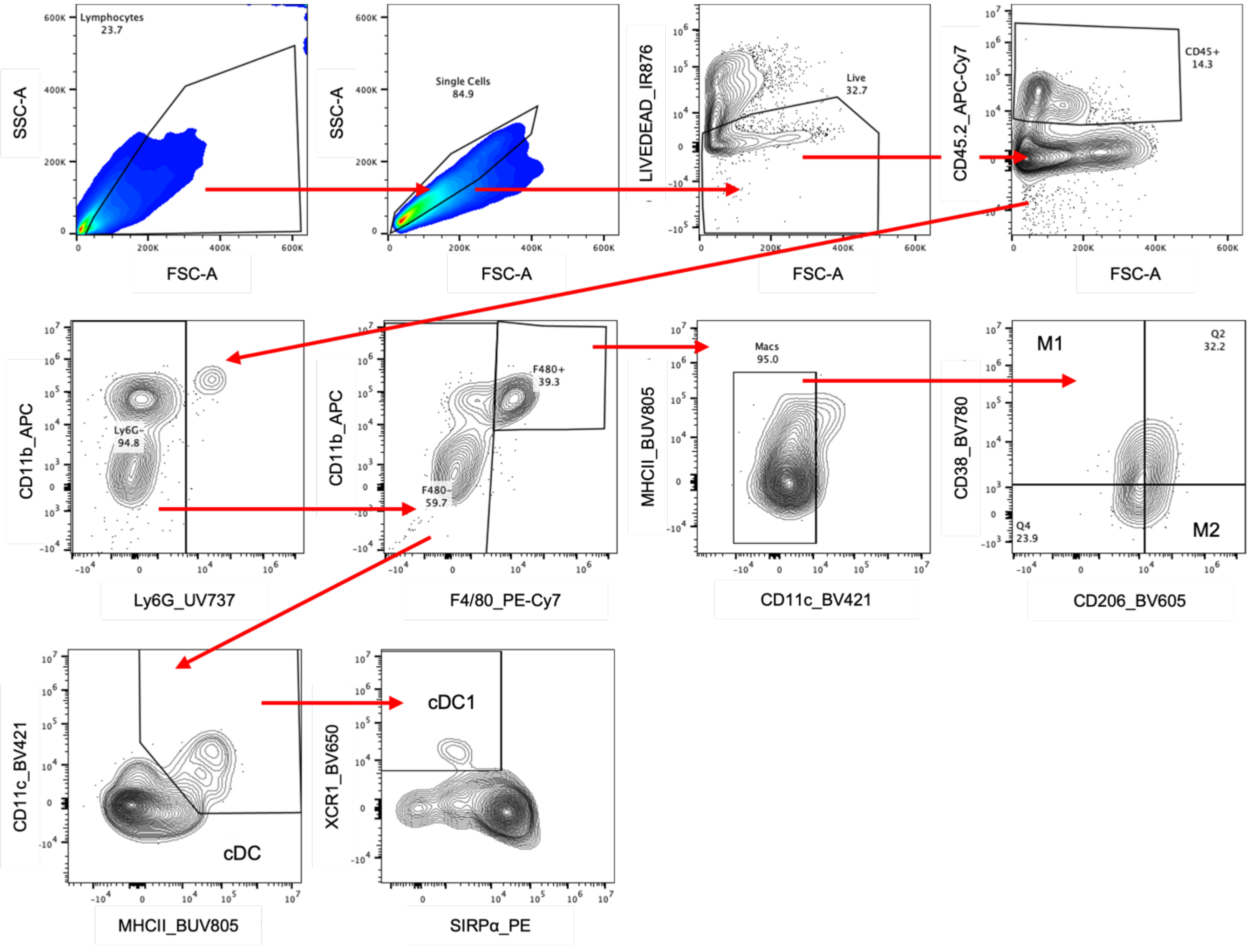


**Figure S5.** Gating strategy for characterizing myeloid cells in the CBD-IL-12 + Dox combo therapy. Cells were first gated based on LIVE/DEAD™ Fixable Near IR (876) Viability Kit followed by staining with indicated markers.
